# Individual selection leads to collective efficiency through coordination

**DOI:** 10.1101/422287

**Authors:** Arthur Bernard, Nicolas Bredeche, Jean-Baptiste André

**Author notes:** Correspondence: Correspondence ✉. J.-B.A. and N.B. contributed equally to this work.

## Abstract

Social interactions involving coordination between individuals are subject to an “evolutionary trap.” Once a suboptimal strategy has evolved, mutants playing an alternative strategy are counterselected because they fail to coordinate with the majority. This creates a detrimental situation from which evolution cannot escape, preventing the evolution of efficient collective behaviours. Here, we study this problem using the framework of evolutionary robotics. We first confirm the existence of an evolutionary trap in a simple setting. We then, however, reveal that evolution can solve this problem in a more realistic setting where individuals need to coordinate with one another. In this setting, robots evolve an ability to adapt plastically their behaviour to one another, as this improves the efficiency of their interaction. This ability has an unintended evolutionary consequence: a genetic mutation affecting one individual’s behaviour also indirectly alters their partner’s behaviour because the two individuals influence one another. As a consequence of this indirect genetic effect, pairs of partners can virtually change strategy together with a single mutation, and the evolutionary barrier between alternative strategies disappears. This finding reveals a general principle that could play a role in nature to smoothen the transition to efficient collective behaviours in all games with multiple equilibriums.

---

The success of a collective action often hinges on the coordinated decisions of several individuals. For instance, carrying out a collective hunt implies that all individuals hunt at the same time, agree on a common prey, and pursue the prey in a coordinated manner. Thus, collective efficiency does not depend solely on the skills of a single individual but emerges from the ability of the group to act together (1–3). This begs the question of how natural selection, which acts on individuals, can shape such collective behaviours.

This problem can be formalised with a specific class of games called “coordination games” To understand, let us consider a situation in which two hunters must coordinate to capture a prey, but have to make a choice between a prey with a high nutritive value and a prey with a low nutritive value. The strategy of choosing the most nutritious prey is evolutionarily stable. If everyone chooses this prey, one’s best response is to choose this prey as well. But choosing the poorly nutritious prey is also evolutionarily stable. If everyone chooses the poorly nutritious prey, there is nothing better one can do than choose the same. The existence of this second, suboptimal, evolutionarily stable strategy (ESS) raises a problem because evolution can hardly move from one ESS to another. If all hunters initially target the low-value prey, mutants preferring the high-value prey are counterselected by frequency-dependent selection because they fail to coordinate with the majority. Hence, individuals are trapped in a suboptimal ESS and collective efficiency is not maximised.

All collective actions where individuals need to coordinate with one another, and where they can do so in either an efficient or an inefficient way, entail such an “evolutionary trap.” Inefficient coordinated behaviours evolve that cannot later be improved by natural selection because *individual* selection has no way of improving *collective* efficiency in a coordination game.

Evolutionary game theoreticians and evolutionary biologists have explored two hypotheses that can explain how this problem can be solved in nature. The first hypothesis is based on stochastic effects (4–6; see also 7 in a different setting). In a finite population fixed in a particular ESS, counterselected mutants can rise in frequency due to genetic drift, and eventually destabilise the existing ESS, thereby moving the population away from the evolutionary trap, toward another, generally superior, ESS. The second hypothesis is based on group selection (8–11). Due to chance, different groups of individuals may initially evolve different ESSes of the same game. If these groups compete with one another, the groups that happen to play the most efficient ESS will eventually prevail, allowing this strategy to spread in the entire population. In sum, according to available theories, coordination games suffer from an evolutionary trap problem, and collective efficiency in these games can ensue either from demographic stochasticity or from group selection, but not from plain individual selection.

However, so far, coordination games have been formally studied in models that were highly stylized, in particular with regard to the mechanistic underpinning of behaviour, and these simplifications may have important consequences. In this paper, we describe simulations of a coordination game using evolutionary robotics (12, 13). As compared to classic evolutionary game-theoretical approaches, evolutionary robotics provides a more realistic modeling of individuals and their environment (14, 15), capturing in particular the practical problems raised by coordination (16).

In this setting, we show that the evolutionary trap actually disappears altogether. Robots evolve a behavioural solution to coordinate with one another that generates indirect genetic effects (17), hereby changing “the rules of the game”. Through their own behaviour, robots transform a coordination game with multiple ESSes and an evolutionary trap problem into a simple individual optimization problem with a single, maximally efficient, ESS. In this transformed game, collective efficiency is reached by plain individual selection, with no need for genetic drift or group selection. We posit that many collective optimization problems may be solved in similar ways in nature.

## Results

We simulate a collective hunt in which two players must coordinate and attack together the same prey to gain a benefit. The two-dimensional environment contains two types of prey, poorly nutritious prey called *boars* (worth 125 payoff units for each individual hunter), and highly nutritious prey called *stags* (worth 250 units for each hunter). Hunting alone is possible, but it provides 0 payoff unit (cf. payoff matrix in Table 1). This game features two Nash equilibriums (and thus two ESSes): to hunt either boars or stags, with the latter equilibrium providing a higher payoff. In technical terms, hunting stags is called the “payoff-dominant” equilibrium.

**Table 1.**
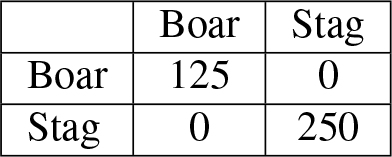
Possible outcome of a two-person coordination game. The game features two hunters. Each player may choose to hunt either a *boar*, or a *stag*. This payoff matrix has two equilibriums, a suboptimal one (both players hunt the boar) and an optimal one (both players hunt the stag).

The robots we use as players are each driven by a multilayer perceptron (18) that maps sensory inputs to motor outputs, with neural weights subject to artificial evolution. Each robot is endowed with proximity sensors all around its body. These sensors are capable, within a limited range, of discriminating between boars, stags, the other robot, and walls. So as to maintain the prey density constant, a captured boar (or stag) is removed from the environment and relocated to a new position (*Methods*).

### Individual selection cannot generate collective efficiency, in a simple setting

The first question we address is whether the evolutionary transition from the least efficient to the most efficient ESS can occur.

First, we let 30 independent populations of robots pre-evolve for 3000 generations with only 1 boar and 1 stag, and modified payoff values: hunting stags temporarily yields no reward. We thus ensure that these 30 populations all evolve the boar-hunting equilibrium, with all individuals always targeting boars and avoiding stags.

Second, each of these 30 populations of evolved boar hunters is used as the seed for another 6000 generations of evolution, with the regular rewards for each prey reinstated (Table 1). In spite of its collective superiority, stag hunting never evolves within the next 6000 generations for any of 30 independent replicates (Figure 1). In every replicate, the mean proportion of stags hunted remains at 0 throughout the 6000 generations. Hence, individuals are genuinely trapped in the suboptimal equilibrium. That is, collective efficiency cannot ensue from plain individual selection.

**Fig. 1.**
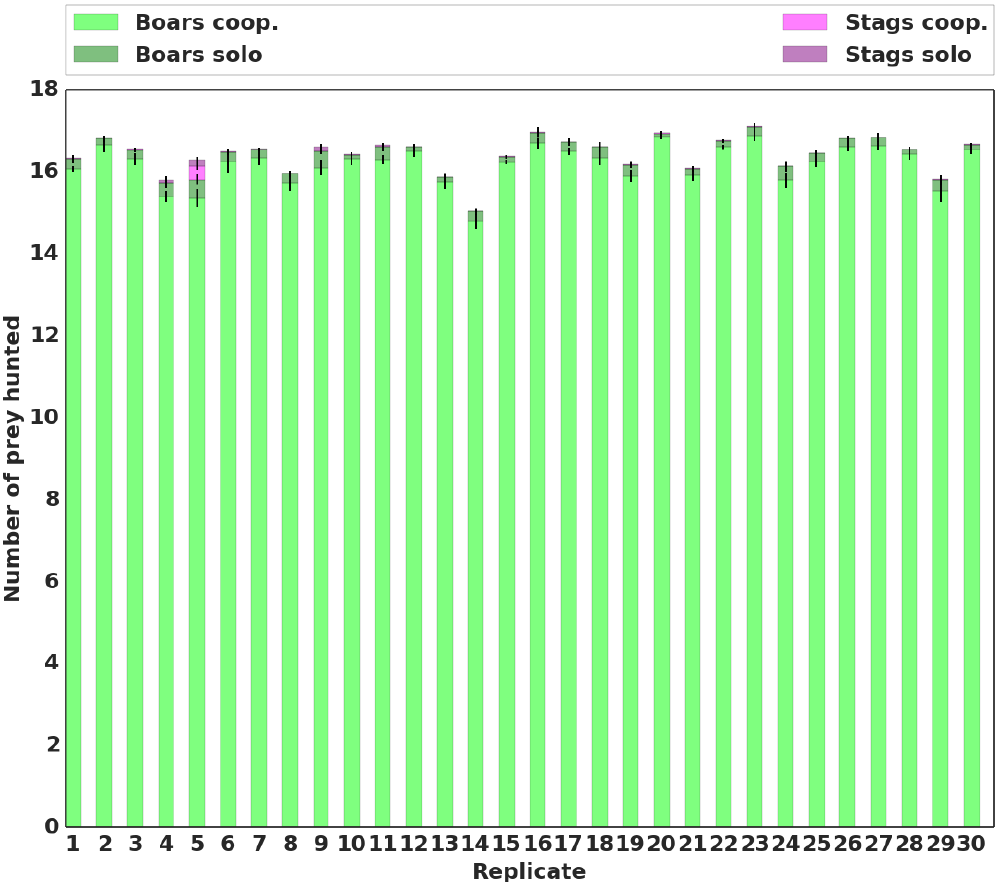
Mean proportion of prey hunted. Repartition of the prey hunted by the best individual in each replicate, at the last generation of evolution (generation 6000). We differentiated between the type of prey hunted (boar or stag). Rewards were 125 for a boar and 250 for a stag (Table 1).

### Collective efficiency can be achieved by individual selection, in a more complex setting

In practice, predators are unlikely to live in a world with a single prey of each kind. In a realistic environment, hunters must agree on a specific *individual* prey to hunt (1–3), and not just on the *type* of prey.

To investigate the consequences of this complication, we follow the same procedure as before. We pre-evolve 30 independent populations of pure boar hunters, using modified payoff values, with the stags never bringing any reward, but this time in an environment with several (9) identical boars and several (9) identical stags present.

We then let each of these 30 populations evolve for another 6000 generations with regular payoff values (Table 1) in the same environment with several boars and stags present (*Methods*).

In this setting, we observe that the transition from boar hunting to stag hunting does occur in 12 replicates out of 30 (Figure 2). This significantly differs from the previous results obtained in a simpler environment (One-tailed MannWhitney U test on the number of replicates where the transition happened *p*-value <0.0001). Environmental complexity promotes the evolutionary transition toward the payoff dominant equilibrium in 40% of the replicates.

**Fig. 2.**
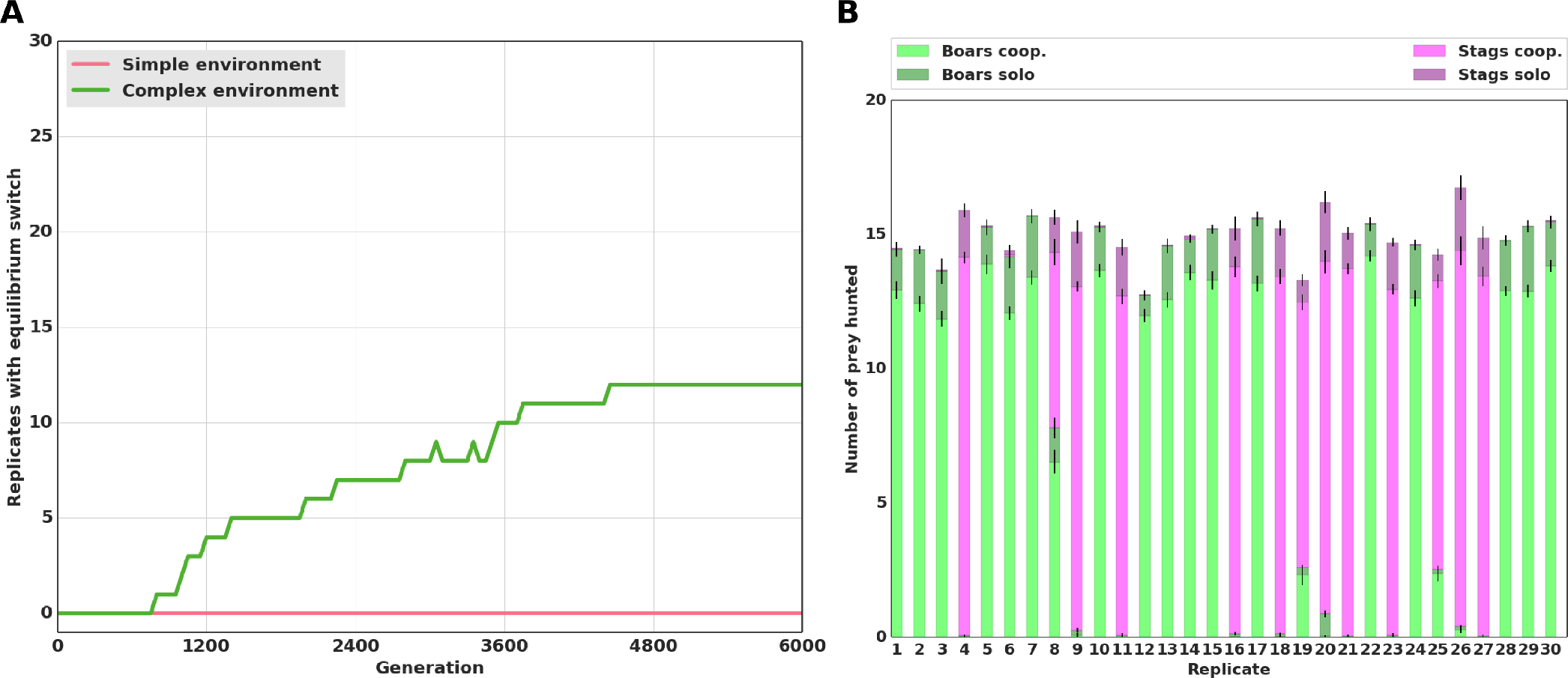
Number of stag-hunting replicates and proportion of prey hunted. *(A)* Number of replicates (out of a total of 30) where stag hunting evolved in the *Simple environment* and *Complex environment* settings. We considered that stag hunting evolved when more than 50% of the prey hunted were stags hunted cooperatively. In the Simple environment setting, the environment was constituted of one boar and one stag. In comparison, in the Complex environment setting, 18 prey were present in the environment, and it was thus necessary to coordinate for cooperation to be possible. *(B)* Repartition of the type of prey hunted by the best individual in every replicate, at the last generation of evolution (generation 6000), in the Complex environment setting. Rewards were 125 for a boar and 250 for a stag (Table 1).

Taking a closer look at these 12 “successful” replicates reveals a particular kind of coordination strategy for collective hunting. Because the environment is more complex, individuals need to react to each other’s behaviour to stay together and converge on the same prey. To this end they evolve a behavioural strategy, which we refer to as the “turning” strategy, whereby they constantly turn around one another. This strategy ensures that they keep their partner in their line of sight and move toward a prey at the same time. Due to their proximity, an individual who gets on a prey is likely to be joined quickly by their partner (Figure 3, a video of this strategy is also available in Supporting Information).

**Fig. 3.**
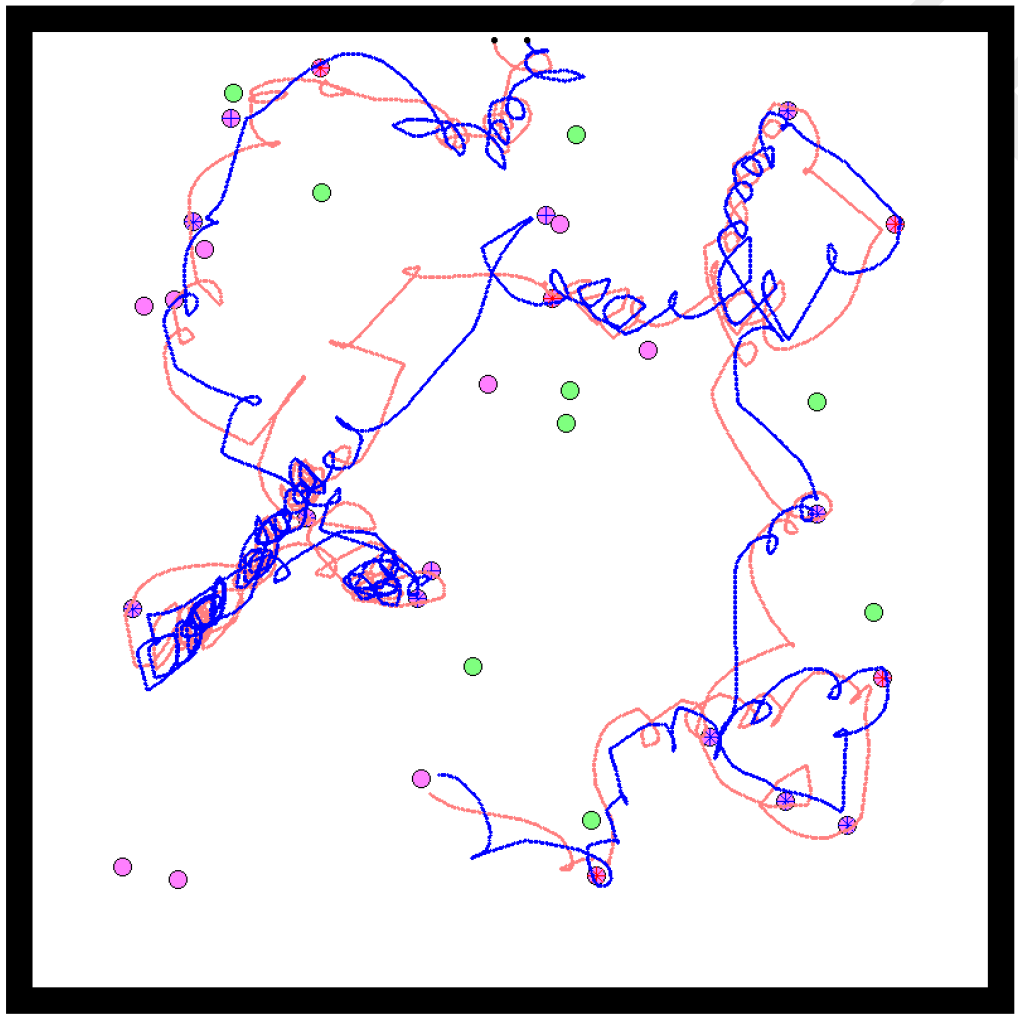
Example of the turning strategy. Example of a simulation where both individuals adopted a *turning* strategy. The path of the agents during this simulation is represented in red and blue, starting from their initial positions (represented by black dots). Each disc represents a prey in the environment. Boars are represented in green and stags in purple. When a prey was killed cooperatively, a red cross (resp. blue) is shown on the prey if the red agent (resp. blue) arrived at this prey first.

This behavioural strategy has an evolutionary implication. Because hunters react to each other’s behaviour, a mutation affecting the behaviour of one individual can also modify the behaviour of her partner. A mutant attracted to stags rather than boars may thus succeed in hunting by transforming, albeit temporarily, her partner into a stag hunter as well. As a result, the evolutionary transition away from the suboptimal trap is facilitated in comparison with the simple environment.

### Division of labour further facilitates collective optimisation

The turning strategy obtained so far results from both hunters demonstrating identical behaviours, whether related to moving together or selecting a prey. While this similarity in behaviour makes it possible for hunters to coordinate toward hunting the same prey, both individuals spend a significant amount of time turning around one another. This results in a tedious process of targeting a particular prey. The question is open as to the existence of more efficient coordination patterns, in particular with respect to assuming complementary behavioural strategies.

We posit that one possible limitation is the lack of expressivity of our choice of control function, a limitation that may not exist in nature. Though multilayered perceptrons are theoretically universal approximators, this is hardly the case in practice (19). In particular, the ability to switch from one behavioural pattern to a completely different one may be required for breaking behavioural symmetry between two individuals, but may be hindered by the limitation of the controller used thus far.

To explore the possible benefits of more complex decisionmaking capabilities, we enable each robot with the possibility to choose from two (possibly very) different controllers, depending on the context at hand. To do so, we introduce a new evolutionary operator: the *network duplication* operator. Loosely inspired by gene duplication (20), a newly created individual may be subject to the complete duplication of its artificial neural network. Conversely, an individual with two networks may have one deleted. In this setup, any individual may possess either one or two network(s). Whenever two individuals who possess two networks each interact with one another, we ensure that each of them expresses a different copy of their own networks (*Methods*).

We use the same experimental procedure as before: 30 populations are pre-evolved independently where only boar hunting yields a reward, and this time we introduce network duplication. We observe a significant difference with respect to previous experiments, as in *all* 30 replicates, we observe the evolution of asymmetrical hunting behaviours in the form of a *leader-follower* division of labour.

Results are also significantly different than in both previous treatments when the original payoff matrix is reinstated (Table 1). The transition to stag hunting occurs in 22 replicates out of 30 (Figure 4), as compared to 0 in the simple environment, and 12 in the complex environment without network duplication (One-tailed Mann-Whitney U test on the number of replicates where the transition happened: *p*-value <0.0001 and *p*-value = 0.015, respectively). As in the pre-evolution runs, the *leader* guides the pair toward a given prey, always arriving first, while the *follower* keeps the leader in its line of sight at all times and joins her afterwards on the prey (Figure 5, a video of this strategy is also available in Supporting Information).

**Fig. 4.**
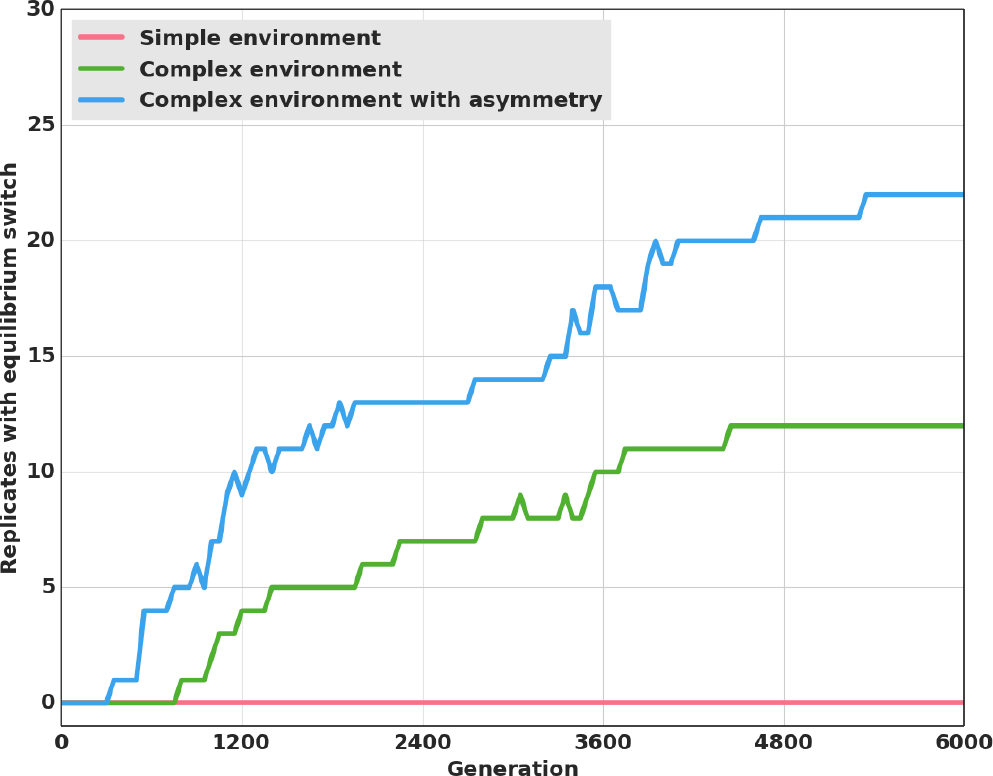
Transitions to the optimal equilibrium through evolutionary time. Number of replicates (out of a total of 30) where the switch to stag hunting occurred in the *Simple environment*, *Complex environment* and *Complex environment with asymmetry* settings. We considered that stag hunting evolved when more than 50% of the prey hunted were stags hunted cooperatively. The Simple environment and Complex environment settings are the same as previously (Figure 2). In the Complex environment with asymmetry, a duplication event could occur where individuals would coevolve two neural networks. We enforced the asymmetry by forcing each hunter to use a different random neural network as its controller. Rewards were 125 for a boar and 250 for a stag (Table 1).

**Fig. 5.**
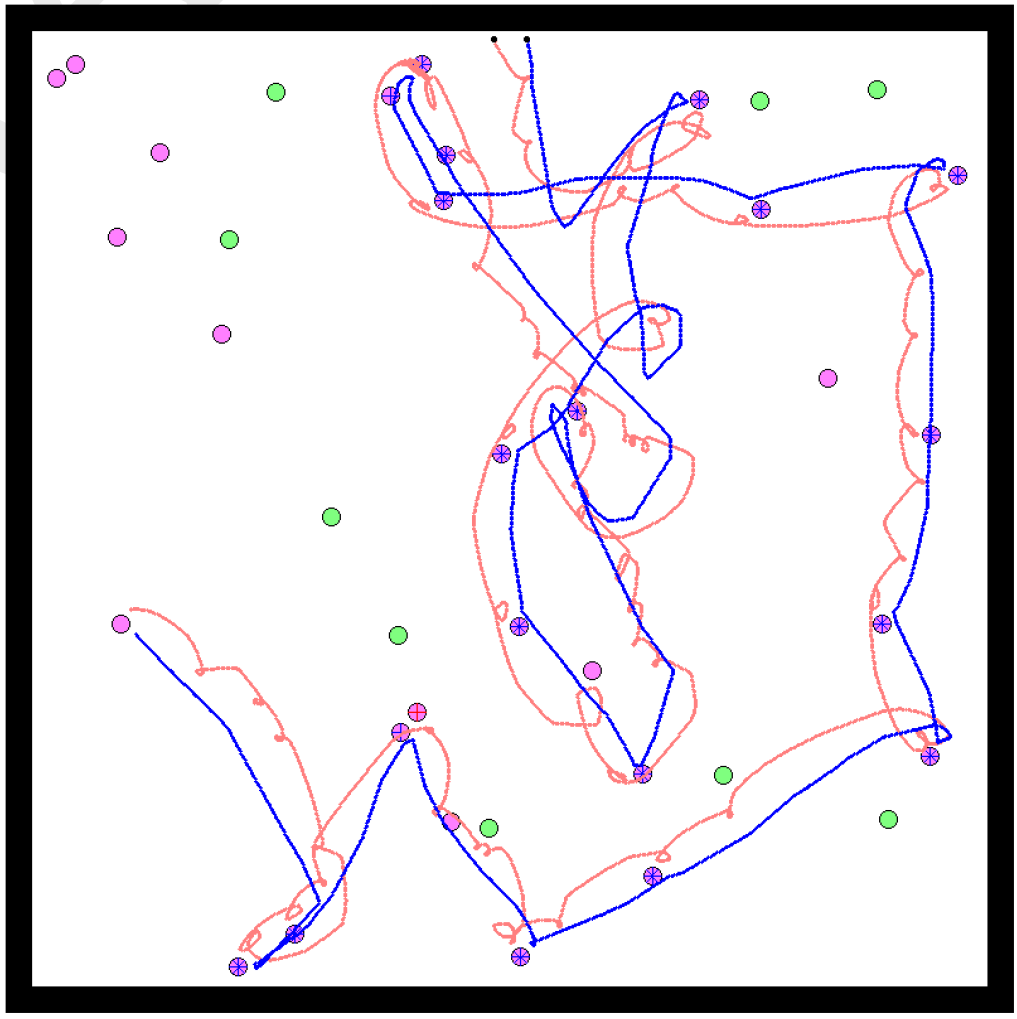
Example of the leader-follower strategy. Example of a simulation where both individuals adopted a *leader-follower* strategy. The path of the agents during this simulation is represented in red and blue, starting from their initial positions (represented by black dots). Each disc represents a prey in the environment. Boars are represented in green and stags in purple. When a prey was killed cooperatively, a red cross (resp. blue) is shown on the prey if the red agent (resp. blue) arrived on this prey first.

This cognitive division of labour stems directly from the duplication of the neural controller: once duplicated, one version of the network always ends up encoding for the leader behaviour, while the other encodes for the follower behaviour, just like duplicated genes of the same family encode slightly different functions.

The division of labour has two consequences. First, it improves hunting efficiency. In the turning strategy, the symmetry of decision-making sometimes hinders the ability to reach a consensus. Even though turning promotes coordination, individuals often still fail to converge on the same prey. In comparison, performance is significantly higher in the leaderfollower strategy (Figure 6, One-tailed Mann-Whitney U test on the mean reward at last generation, *p*-value <0.001): the frequency of coordination failures is reduced thanks to a clear separation of roles.

**Fig. 6.**
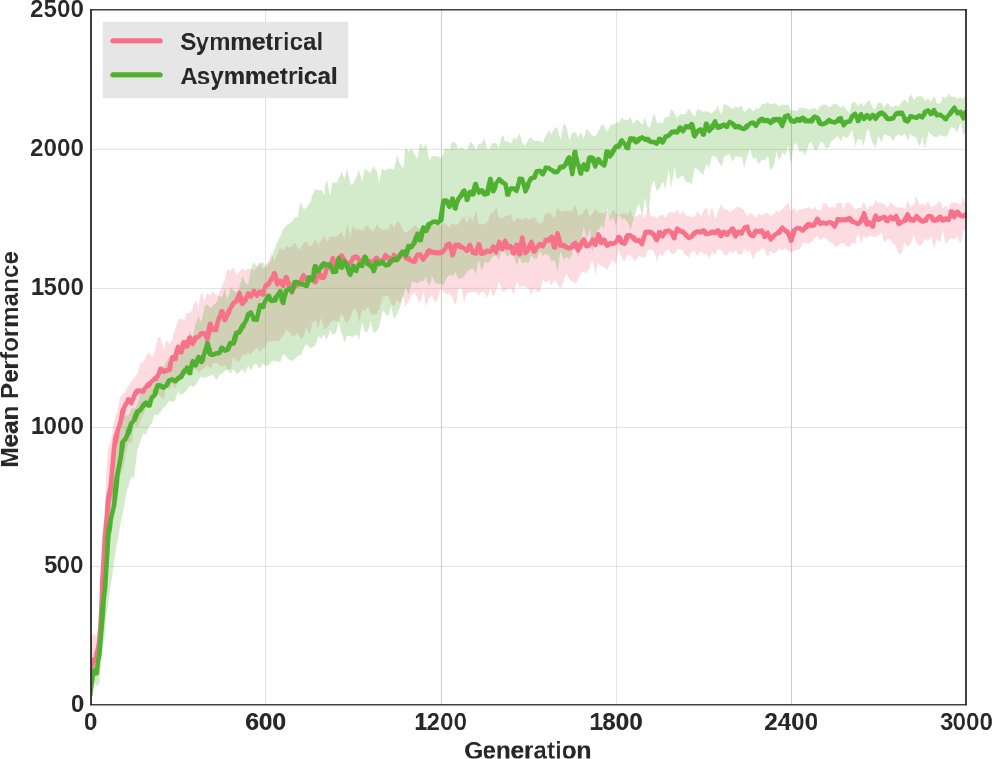
Mean reward comparison of the turning strategy and the leaderfollower strategy. Mean reward over evolutionary time of the best individuals in each of the 30 replicates when adopting a turning strategy or a leader-follower strategy during the pre-evolution step (which lasted for 3000 generations). Rewards were 125 for a boar and 250 for a stag (Table 1).

Second, the division of labour has an evolutionary consequence. In the leader-follower strategy, just as in the turning strategy, hunters react to each other’s behaviour and are therefore also prone to react to mutants’ behaviour. But, in contrast to the turning strategy, this response is asymmetrical and, therefore, more precise. Any mutation affecting the leader’s behaviour also changes completely the behaviour of the follower. That is, a mutation in a single individual automatically affects two individuals at the same time. As a consequence, the adaptive valley between boar hunting and stag hunting disappears or, put differently, boar hunting ceases to be an equilibrium. Any increase in the probability of hunting a stag rather than a boar, when playing the role of a leader, is directly favoured by individual selection, and pure stag hunting therefore becomes the only evolutionary equilibrium.

## Discussion

Collective actions often require several individuals to make coordinated choices. As a result, their efficiency, or lack of efficiency, is a collective property, not a property of any particular individual. This raises an evolutionary difficulty because natural selection acts on individual, not collective, properties. Collective actions are thus subject to an “evolutionary trap” problem. Once a relatively successful but still perfectible collective organisation has evolved, any single mutant playing a better strategy will be counterselected due to her lack of coordination with others. For collective efficiency to be reached by evolution, several individuals would all somehow have to “mutate collectively”, but genetic mutations do not occur in several organisms at the same time.

In this paper, we studied this problem in artificial robotics experiments. We simulated the life and the long-term evolution of a population of simple robots that played a 2 x 2 coordination game. Robots were hunters who could choose between two types of prey that were either poorly nutritious or highly nutritious. But they could only be successful if they converged together on the same prey. Hence, they faced a coordination problem with two ESSes - hunting poorly nutritious prey or hunting highly nutritious prey, and an adaptive valley in between. Our aim was to find out how the evolutionary trap problem materialises and how it is solved - or not solvedin a model possessing a greater degree of realism than conventional models of coordination games.

We first confirmed the existence of an evolutionary trap. In a simple setting where the environment was constituted of two individual prey only - one poorly nutritious and one highly nutritiousif we initially forced the robots to play the suboptimal ESS (attacking the poorly nutritious prey), all populations of robots remained stuck in this ESS “forever,” that is, at least for the 6,000 generations of our simulations. Individual mutants who were targeting the better prey could not be favoured due to their singularity.

This observation may seem at odds with evolutionary game-theoretical “drift” models (4–6). According to these models, finite populations always eventually escape from evolutionary traps, because counterselected mutants rise in frequency by genetic drift and eventually replace the suboptimal resident. Mathematical analyses of this process show that, in the long run, populations should spend most of the time in the vicinity of one specific equilibrium (called the “stochastically stable” equilibrium), which corresponds to stag hunting with our parameter setting. Hence, “drift” models predict that our populations should not be trapped in the suboptimal strategy.

This discrepancy is a matter of time scale, however. According to drift models, our robots should eventually escape from the evolutionary trap and hunt stags, in the “long” run, but the question is how “long” in practice? The answer to that question depends a great deal on the practical availability of mutants. In game-theoretical models, stag hunters are simply assumed to occur by mutation from boar hunters at a given rate. In a robotic setting like this one, however, stag hunters must appear by random changes in the connection weights of boar hunters’ neural networks, and multiple such changes separate a pure and well-optimised boar hunter from a pure and well-optimised stag hunter. As a result, mutants playing the stag-hunt strategy are extremely rare in a population of pure boar hunters.

This is confirmed in a supplementary experiment (see Fig. 7, and *Methods*), where we analysed the behaviour of 105 mutants generated randomly from a pure and well-optimised “boar hunter” genotype. This analysis showed that, at best, the random mutants merely had a probabilistic tendency to target the stag. That is, they played a mixed rather than a pure strategy. These intermediate mutants can never prevail in a population of pure boar hunters, even after a phase of genetic drift, because mixed strategies are strongly counterselected in coordination games (owing to the uncertainty they generate). Hence, these mutants cannot bring about the stochastic transition to stag hunting.

**Fig. 7.**
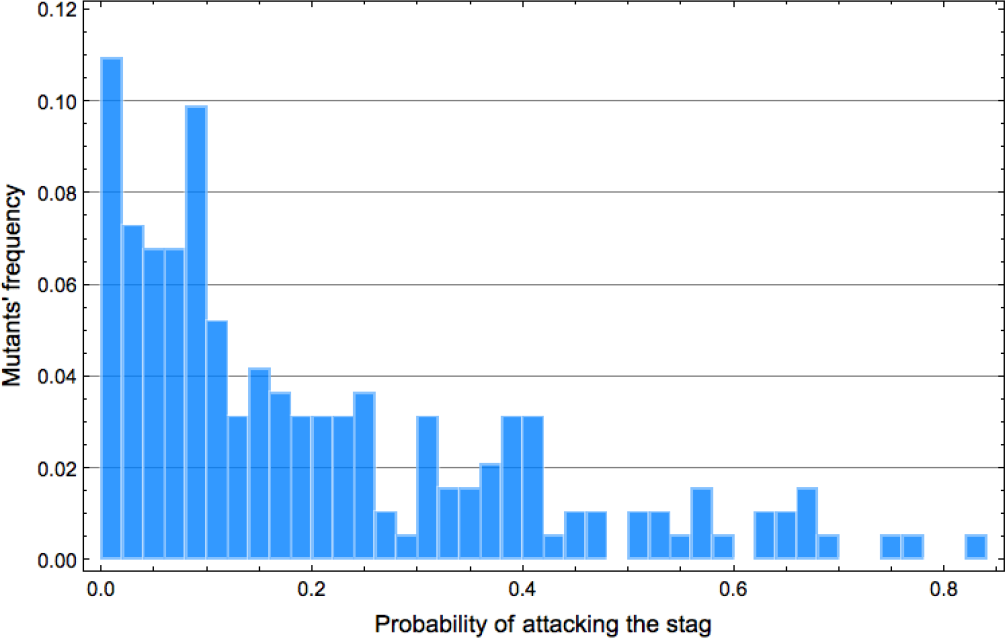
Probability to prefer stags over boars in mutants generated randomly from a pure boar-hunter genotype. Among the 105 mutants generated, we display only the 192 mutants whose probability to hunt stags is greater than 1% (see *Methods*).

Although the occurrence of strong-effect mutants able to destabilise the suboptimal equilibrium is possible in principle, the above analysis shows that it is highly unlikely in practice owing to the mutational distance between ESSes. As a result, the stochastic evasion from the evolutionary trap is an extraordinarily slow process in our simulations. Because the mutational distance between ESSes will often be even greater in biology than in the present setting, stochastic evasions from suboptimal equilibriums are presumably a highly improbable event in general in biological settings.

However, we then showed that this problem actually disappears in a more realistic setting. In a richer environment constituted of several prey of each kind - several poorly nutritious prey and several highly nutritious prey, individuals needed to actively coordinate with their partner to converge on the same prey. To resolve this problem, they evolved behavioural tactics to keep track of and follow their partner. In our experiments we observed the evolution of two such tactics. In the first series of experiments, individuals constantly turned around one another, never moving away from their partner, which increased the probability that they both would eventually converge on a prey. In other experiments where we authorised a behavioural asymmetry between partners, individuals evolved a leader/follower strategy whereby a single individual chose a prey, whereas the other simply followed her.

These coordination strategies evolved because they had immediate individual benefits. They increased the probability for individuals to hunt successfully. But they also had an unintended evolutionary consequence. When individuals had the capacity to coordinate with each other, a mutation affecting the behaviour of one individual also indirectly modified, phenotypically, the behaviour of his/her partner, almost as if individuals had mutated “collectively.” In quantitative genetics, such an effect is called an “indirect genetic effect” (17) because a gene affects the phenotype of an individual in which it is not directly expressed. Indirect genetic effects are well known for changing the evolutionary process in sometimes dramatic ways, by altering the genotypephenotype relationship. In the present case, behavioural coordination tactics evolved to deal with the uncertainty of the “normal” environment in which one’s partners only targeted suboptimal prey but the precise individual prey they were targeting could vary, which required being able to follow them. However, once evolved, coordination tactics also happened to work efficiently when interacting with mutants who preferentially targeted other types of prey. They led one to follow and coordinate with mutants like they did with “normal” residents. Consequently, genetic mutants that preferred targeting the most nutritious type of prey were directly favoured by individual selection because they always had a resident who accepted to follow them since she was indirectly influenced by their mutated gene. The suboptimal coordination equilibrium was no longer an evolutionary trap. In finding a solution to the *behavioural* coordination problem, individuals solved the *evolutionary* coordination problem as well.

Put another way, behavioural coordination strategies changed the nature of the game. Individuals initially played a coordination game in which two players needed to jointly evolve a compatible preference. This raised a bootstrapping problem and made the transition from one equilibrium to another unlikely. By evolving endogenously a coordination strategy, individuals turned this game into a plain optimization game in which a single player was simply selected to choose the best possible prey.

Beyond the particular setting considered in this paper, we think these results reveal a general principle that could play a role in all games with multiple equilibriums, that is, in all coordination games, but also in repeated games such as the repeated prisoner’s dilemma (see, for instance, 21, 22). As a rule, there are many reasons why the behaviour of one’s partners will vary in all these games, making it necessary for one to adapt plastically to this variability (23). In our simulations, for instance, individuals evolved a coordination strategy to adapt to the precise location where their partner was heading, but the same principle should hold in other settings as well. Even though behavioural plasticity originally evolves merely to deal with partners’ phenotypic variability, it also happens to generate an adaptive response in front of mutants. These mutants probably did not exist when behavioural plasticity evolved, but they nevertheless happen to trigger the exact same response. And because this response was originally meant to maximise efficiency, it is likely to do so with mutants, too, as our experiments illustrate. Hence, there is a general reason why the plastic response of individuals to each other should often “change the rules of the game” and smoothen the transition to efficient collective behaviours.

## Methods

### Experimental setup

The environment is a 800 by 800 unit arena with four solid walls. Each simulation is conducted with a pair of hunters (the robotic agents) and a varying number of prey of two types, boars and stags with respective rewards 125 and 250 (Table 1). The initial positions of the prey are random and the prey cannot move. To capture a prey, the two hunters have to stay in contact with it for 800 time steps (out of a total of 20000 time steps for each simulation). Both robots have to be in contact with the prey at the end of the 800 time steps for the hunt to be considered successful. Once captured, the prey is removed and replaced at a random position in the arena. In the “simple” environment condition, there is always exactly one boar and one stag present in the environment. In the “complex” environment condition, there is always 9 prey of each type. Robots (that is, hunters) begin the simulation next to each other at the top of the arena and can then move freely in the environment. To do so, they are equipped with a set of sensors and two independent wheels connected by a fully connected multilayer perceptron. Sensors comprise 12 proximity sensors and a camera. Proximity sensors are evenly distributed around the robot’s body, and each has a range of 40 units. A proximity sensor is a ray toward a particular direction indicating to the robot the distance of the first obstacle in this direction. The camera is placed on the front of the robot, and its 90 degree field of view is divided into 12 equally spaced rays. Each ray of the camera indicates the type (that is, hunter, boar, or stag) and the proximity of the nearest agent in its direction. Robots are individually controlled by a fully connected multilayer perceptron with a single hidden layer. The inputs of the neural network are fed with the sensory data of the robot. One input neuron is used for each of the 12 proximity sensors, with maximal (resp. minimal) neural activity when the agent is directly in contact with an obstacle (resp. when there is no obstacle in the range of the sensor). Three neurons are used for each of the 12 rays of the camera: two neurons to encode the type of obstacle in a two-bit binary value and one neuron to encode the proximity of the obstacle. Finally, there is a bias neuron whose value is always equal to one. The total number of input neurons is 49. The hidden layer contains 8 neurons, while the output layer contains 2 neurons. These 2 output neurons control the speed of the left and right wheels; minimal (resp. maximal) activity results in maximal backward (resp. forward) actuation. The activation function used to compute outputs is a sigmoid function. Connection weights are each encoded in a single gene (the total genome size is 410).

### Simulating artificial evolution

In each of the 30 independent replicates, we let a population of 20 individuals evolve. Each individual is encoded as a genome, where each gene codes for a connection weight of the multilayer perceptron controller. Every gene in the genome is first initialised with a random value sampled uniformly in [0, 1]. In each generation, the performance of every individual is evaluated by matching her with five different random partners. In turn, the performance of each pair of partners is evaluated through five independent trials. Hence, the fitness of every individual is computed in each generation as an average across 25 independent trials. We then apply a (10 + 10) elitist selection strategy (24). Generation *t* + 1 is composed of the 10 best individuals of generation *t* plus 10 mutants generated from a single parent of generation *t*. Mutations are sampled according to a Gaussian operator, with a standard deviation of 2 × 10^−1^ and a per-gene mutation probability of 5 × 10^−3^.

### Duplication and coevolution of neural networks

To study the effect of an asymmetry between hunters, we allow the duplication of neural networks. Every individual initially has a single neural network but duplication and deletion events can occur randomly (at the same moment of the life cycle than mutation). When duplication occurs, each gene is duplicated to create a new genome encoding for a second neural network that can then evolve independently of the first. When deletion occurs, one of the two neural networks of the individual is deleted randomly. Duplication occurs with a probability 5 × 10^−2^ and deletion with a probability 5 × 10^−3^ per generation.

### Analyses of boar-hunter mutants w.r.t. stag hunting

We generate 100.000 random mutants from a well-optimised boar-hunter genotype (with the same mutation parameters than in our evolutionary simulations), and assess each mutant’s hunting preferences. From among these 100.000 mutants, we extract 192 mutants that displayed a probability greater than 0.01 of hunting the stag. Figure 7 shows the distribution of the preferences of these 192 mutants. Most mutants have only a small probability to hunt stags. In particular, not a single pure stag hunter can be found among the 100.00 mutants.

## Data availability

The datasets generated and analysed during the current study are available at http://pages.isir.upmc.fr/~bredeche/data/PaperOnIndividualSelection_data_2018.tar.gz

## Code availability

The source code of all simulations presented in the current study is available on GitHub: https://github.com/CAThanatos/sferes-leaderfollower

## ACKNOWLEDGEMENTS

This work is supported by the European Union’s Horizon 2020 research and innovation program under grant agreement No 640891 (DREAM project). Experiments presented in this paper were carried out using the Grid’5000 experimental

## Bibliography

1. Michael S Alvard and David A Nolin. Rousseau’s Whale Hunt? 43(4):533–559, 2002.

2. Michael S Alvard. Kinship, Lineage, and an Evolutionary Perspective on Cooperative Hunting Groups in Indonesia. Human Nature, 14(2):129–163, 2003.

3. Christine M. Drea and Allisa N. Carter. Cooperative problem solving in a social carnivore. Animal Behaviour, 78(4):967–977, oct 2009. ISSN 00033472. doi: 10.1016/j.anbehav.2009.06.030.

4. Dean Foster and Peyton H Young. Stochastic Evolutionary Game Dynamics. Theoretical Population Biology, 38:219–232, 1990.

5. Michihiro Kandori, GJ Mailath, and Rafael Rob. Learning, Mutation, and Long Run Equilibria in Games. Econometrica, 61(1):29–56, 1993.

6. H. Peyton Young. The Evolution of Conventions. Econometrica, 61(1):57, 1993. ISSN 00129682. doi: 10.2307/2951778.

7. M A Nowak, A Sasaki, C Taylor, and D Fudenberg. Emergence of cooperation and evolutionary stability in finite populations. Nature, 428(6983):646–650, 2004.

8. R Boyd and P J Richerson. Group Selection among Alternative Evolutionarily Stable Strategies. Journal of Theoretical Biology, 145(3):331–342, 1990.

9. R Boyd and P J Richerson. Group beneficial norms can spread rapidly in a structured population. J Theor Biol, 215(3):287–296, 2002.

10. Peter J Richerson and Robert Boyd. Not by genes alone : how culture transformed human evolution. University of Chicago Press, Chicago, 2005.

11. Robert Boyd and Peter J Richerson. Voting with your feet: Payoff biased migration and the evolution of group beneficial behavior. Journal of Theoretical Biology, 257(2):331–339, 2009. ISSN 00225193. doi: 10.1016/j.jtbi.2008.12.007.

12. Stefano Nolfi and Dario Floreano. Evolutionary Robotics: The Biology, Intelligence, and Technology of Self-Organizing Machines. MIT Press, 2000. ISBN 0262140705.

13. S. Doncieux, N. Bredeche, J.-B. Mouret, and A.E. Eiben. Evolutionary robotics: what, why, and where to. Frontiers in Robotics and AI, 2, 2015. doi: 10.3389/frobt.2015.00004.

14. Sara Mitri, Steffen Wischmann, Dario Floreano, and Laurent Keller. Using robots to understand social behaviour. Biological reviews of the Cambridge Philosophical Society, 88(1): 31–9, feb 2013. ISSN 1469-185X. doi: 10.1111/j.1469-185X.2012.00236.x.

15. Vito Trianni. Evolutionary Robotics: Model or Design? Frontiers in Robotics and AI, 1 (December):1–6, dec 2014. ISSN 2296-9144. doi: 10.3389/frobt.2014.00013.

16. Arthur Bernard, Jean-Baptiste André, and Nicolas Bredeche. To cooperate or not to cooperate: Why behavioural mechanisms matter. PLoS Comput Biol, 12(5):1–14, 05 2016. doi: 10.1371/journal.pcbi.1004886.

17. Jason B. Wolf, Edmund D. Brodie III, James M. Cheverud, Allen J. Moore, and Michael J. Wade. Evolutionary consequences of indirect genetic effects. Trends in Ecology and Evolution, 13(2):64–69, 1998. ISSN 0169-5347. doi: https://doi.org/10.1016/S0169-5347(97)01233-0.

18. D. E. Rumelhart, G. E. Hinton, and R. J. Williams. Parallel distributed processing: Explorations in the microstructure of cognition, vol. 1. chapter Learning Internal Representations by Error Propagation, pages 318–362. MIT Press, Cambridge, MA, USA, 1986. ISBN 0-262-68053-X.

19. G. Cybenko. Approximation by superpositions of a sigmoidal function. Mathematics of Control, Signals and Systems, 2(4):303–314, Dec 1989. ISSN 1435-568X. doi: 10.1007/BF02551274.

20. Austin L Hughes. Adaptive evolution after gene duplication. Trends in Genetics, 18(9): 433–434, 2002.

21. R J Aumann and L S Shapley. Long-Term Competition: A game-theoretic analysis. In N Megiddo, editor, Essays in Game Theory in Honor of Michael Maschler. Springer, New York, 1994.

22. R Boyd. Reciprocity: You have to think different, 2006. ISSN 1010061X.

23. John M Mcnamara, Olof Leimar, and Phil Trans R Soc B. Variation and the response to variation as a basis for successful cooperation. Philosophical Transactions of the Royal Society B-Biological Sciences, 365(1553):2627–2633, 2010. doi: DOI10.1098/rstb.2010.0159.

24. David E. Goldberg. Genetic Algorithms in Search, Optimization, and Machine Learning. 1989. ISBN 0201157675. doi: 10.1007/BF01920603.

